# Brain Dynamics of Spatial Reference Frame Proclivity in Active Navigation

**DOI:** 10.1101/2021.02.28.433279

**Authors:** Che-Sheng Yang, Jia Liu, Avinash Kumar Singh, Kuan-Chih Huang, Chin-Teng Lin

## Abstract

Recent research into navigation strategy of different spatial reference frame proclivities (RFPs) has revealed that the parietal cortex plays an important role in processing allocentric information to provide a translation function between egocentric and allocentric spatial reference frames. However, most studies merely focused on a passive experimental environment, which is not truly representative of our daily spatial learning/navigation tasks. This study investigated the factor associated with brain dynamics that causes people to switch their preferred spatial strategy in different environments in virtual reality (VR) based active navigation task to bridge the gap. High-resolution electroencephalography (EEG) signals were recorded to monitor spectral perturbations on transitions between egocentric and allocentric frames during a path integration task. Our brain dynamics results showed navigation involved areas including the parietal cortex with modulation in the alpha band, the occipital cortex with beta and low gamma band perturbations, and the frontal cortex with theta perturbation. Differences were found between two different turning-angle paths in the alpha band in parietal cluster event-related spectral perturbations (ERSPs). In small turning-angle paths, allocentric participants showed stronger alpha desynchronization than egocentric participants; in large turning-angle paths, participants for two reference frames had a smaller difference in the alpha frequency band. Behavior results of homing errors also corresponded to brain dynamic results, indicating that a larger angle path caused the allocentric to have a higher tendency to become egocentric navigators in the active navigation environment.

## I. Introduction

In our daily life, spatial navigation is a complex cognitive task frequently occurring during environment exploration. Spatial representations construct the basis for integrating allocentric and egocentric information while navigating in the environment. As per Klatzky et al. 1998 [1], two distinct types of spatial representations that can be described, one representing entities in space based on an allocentric coordinate system and the other based on an egocentric coordinate system. The system of allocentric representation is an environment-centered or object-centered system that represents the position of one object or navigator with respect to an aspect of other external entities such as landmarks. In the egocentric reference system, the spatial location of an object is determined with respect to the aspect of the navigator or observer. The spatial representation of an object depends on the observer’s position or orientation, and the representation will transform when the observer’s position or orientation changes. In summary, the spatial representation of an object is body-based in the egocentric reference system, but it is object-based in the allocentric reference system [1, 2].

Moreover, people who use allocentric reference frames known as nonturners, while people who use egocentric reference frames known as turners. These categories of people are based on whether their mental map would turn after actual turn during spatial navigation or not [3]. Mental map in nonturners do not turn after turning in actual spatial navigation, and turners turn their mental map. Both allocentric and egocentric spatial reference frames signify different information; spatial navigation requires and is based on the parallel use of egocentric and allocentric representations [4].

Many navigation studies have demonstrated the hippocampus, parahippocampal gyrus, posterior cingulate gyrus, anterior frontal cortex, motor cortex, superior and inferior parietal cortex, occipital cortex, retrosplenial cortex, and temporal cortex are related to spatial information processing [4–6]. Previous studies have found that navigators constantly reveal a preference, indicating that people use only one specific reference frame or a subset of reference frames in various navigation environments [3, 6–11]. However, the phenomenon of switching spatial strategies was revealed by Gramann et al. in 2012 [12] and Ehinger et al. in 2014 [13]. Especially, in Gramann’s study, 39% of participants unexpectedly switched to their non-preferred (allocentric) reference frame for vertical direction changes.

Well-controlled studies under restricted laboratory conditions have contributed enormously to the knowledge about brain processes over the past decades. It remains to be tested whether these assumptions hold and to what degree the results obtained in reduced experimental setups transfer to natural conditions. Specifically, such controlled settings often imply sitting in front of a computer monitor, thus omitting important sensory information that would otherwise be present in a natural environment.

To build up the knowledge between spatial learning/navigation and a more natural experimental environment, the fully immersive virtual reality (VR) protocols are widely applied [14–18]. The immersive VR technology allows a user to actively and naturally explore and sense the pervasive computing environment with the stimulation of visual, auditory, and proprioceptive modalities in combination with high-density EEG [15, 19, 20]. Delaux and his colleagues studied simultaneous brain/body imaging during navigation in mobile conditions with virtual Y-maze by using the VR environment [15]. For spatial reference frames on diverse conditions of spatial navigation, Moraresku and Vlcek investigated brain activation during different reference frame proclivities (RFPs) used in previous VR environment-based studies [21]. Moreover, using a novel, fully mobile virtual reality paradigm, Plank and his colleagues studied the EEG correlates of spatial RFPs formed during unsupervised exploration [14]. There is no doubt that the technical developments of VR unlocked the possibility to investigate spatial navigation in more naturalistic environments with well-controlled experimental parameters.

However, though more research on spatial learning started the investigation in a condition closer to a real-life environment, it remains unclear how switching in spatial strategy work in an active environment. As mentioned in Gramann et al. 2012, ‘s about ~50% of participants tend to switch from preferred RFP to their non-preferred RFPs during navigation tasks [12]. Based on previous literature, some participants seem to be impaired in using the non-preferred RFP when the environment requires them to do so [7, 9]. The preference of RFPs seems to critically depend on individual abilities and experience in different environments [12]. Unfortunately, less research yet has been conducted investigating the switch of RFPs in an active environment. Thus, to investigate individual spatial strategies for navigation, it is crucial to understand the possible factors and conditions that may causes people to switch their preferred spatial strategy in different navigational environments.

To narrow the gap on the lack of RFPs studies under different environment conditions, in our experiment, we built an active environment closely resembling a real walk compared with passive tasks to investigate the performance change and human brain dynamics of the two strategies of spatial reference frames. To do so, we have studied if turning angle is the possible factor behind the change in RFP. Our contributions are as follows:

1. An immersive VR-based active navigation environment to understand and investigate RFPs.
2. An EEG-based biomarker representing a difference in active and passive navigation in the participant.
3. An EEG-based biomarker representing the difference between RFPs among participants.

## II. Materials and methods

### A. Participants

In this study, EEG data were recorded from 21 right-handed male participants in the active environment and 20 right-handed male participants in the passive environment; these were the experimental and control groups, respectively. The mean age was 23.6 years (in a range of 22-27 years) with no prior experience pertaining to the experiment. Following an explanation of the experimental procedure, all participants provided informed consent before participating in the study. This study obtained the approval of the institute’s Human Research Ethics Committee of National Chiao Tung University, Hsinchu, Taiwan. None of the participants were aware of the experimental hypothesis and did not have a history of psychological disorder, which might have affected the experimental results.

### B. EEG Setup

During the EEG-based experiment, EEG signals were recorded using an EEG cap with 32 Ag/AgCl electrodes, referenced to linked mastoids. The placement of the EEG electrodes was consistent with the extended 10% system [22]. The contact impedance between the electrode and the scalp was kept below five kΩ. Before the experiment, channel locations were digitized with a 3D digitizer (POLHEMUS 3 space Fastrak). EEG data were recorded and amplified with the synAmps RT 64-channel Amplifier.

### C. Pretask

This task was designed to distinguish participant’s preferred RFP. We used the Gramann et al. 2010. the 2010 [4] based experiment scenario to evaluate participant’s allocentric and egocentric spatial RFP. In the pretask, participants sat on a chair before the screen and watched a turning-tunnel video based on the script. In the tunnel, participants would see one part that randomly turned left or right before the endpoint. After going through the tunnel, two arrows pointing in two different directions were presented, and participants needed to select the arrow they think is pointing in the direction of the starting point. Fig. 1 shows the turning tunnel and the two arrows that the participants needed to choose between after reaching the end point. If the tunnel turned right, then nonturners would select the arrow on the right side, and turners would select the other one. Participants were to maintain at least 80% accuracy 20 times for one reference frame before beginning the navigation task.

**Fig. 1.**
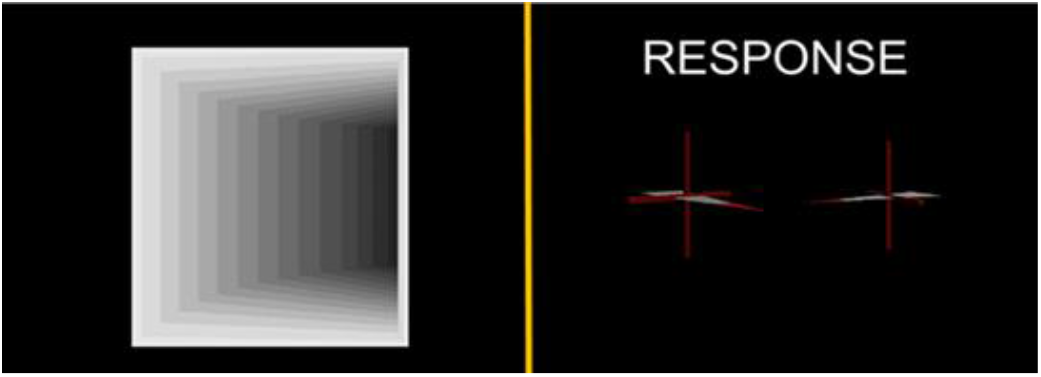
Illustration of the pretask design. Participants would pass through the tunnel, which turned right or left randomly and were to select the arrow pointing toward the start point once they reached the end of the tunnel.

### D. Experimental Environment and Procedure

Participants performed the active experimental task while standing on an Omni treadmill^1^ in the range of two VR base stations and held one HTC controller in their dominant hand (see Fig. 3A). Participants wore an EEG electrode cap and HTC VR head-mounted display that used an OLED display with a resolution of 2160 x 1200 and a refresh rate of 90 Hz. An assistant helped the participants put on the EEG cap first and then put the head-mounted display on. We directly put the top belt of the HTC Vive on top of the central channel of the EEG cap and adjusted it manually to avoid or reduce the pressure applied by the EEG channels. synAmps was fixed on the top to prevent wires from winding around participants during the experiment. (See Fig. 2.) In passive navigation, participants in the control group also wore the EEG electrode cap but did not wear a head-mounted display; they sat in front of the computer screen, watched the scenario, and performed tasks.

**Fig. 2.**
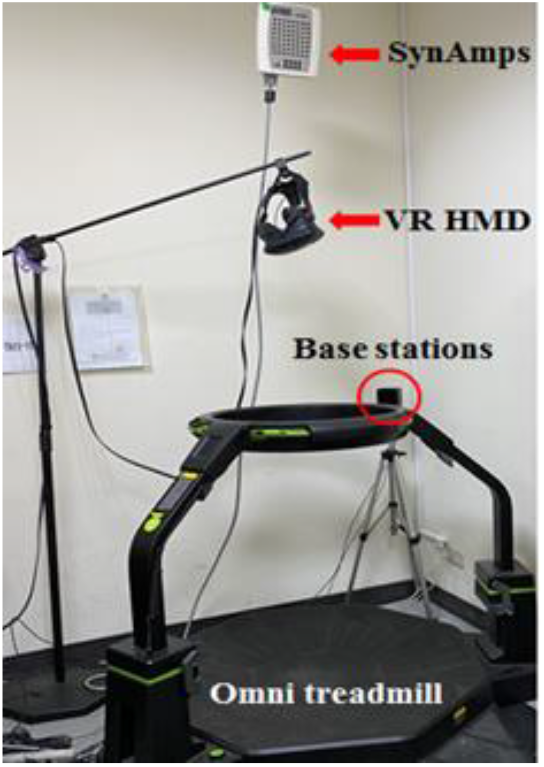
Experiment environment. Participants performed the experiment on a treadmill between two VR base stations that created the range of VR scenarios. SynAmps and VR head-mounted displays were fixed on the top to prevent wires winding around participants.

**Fig. 3.**
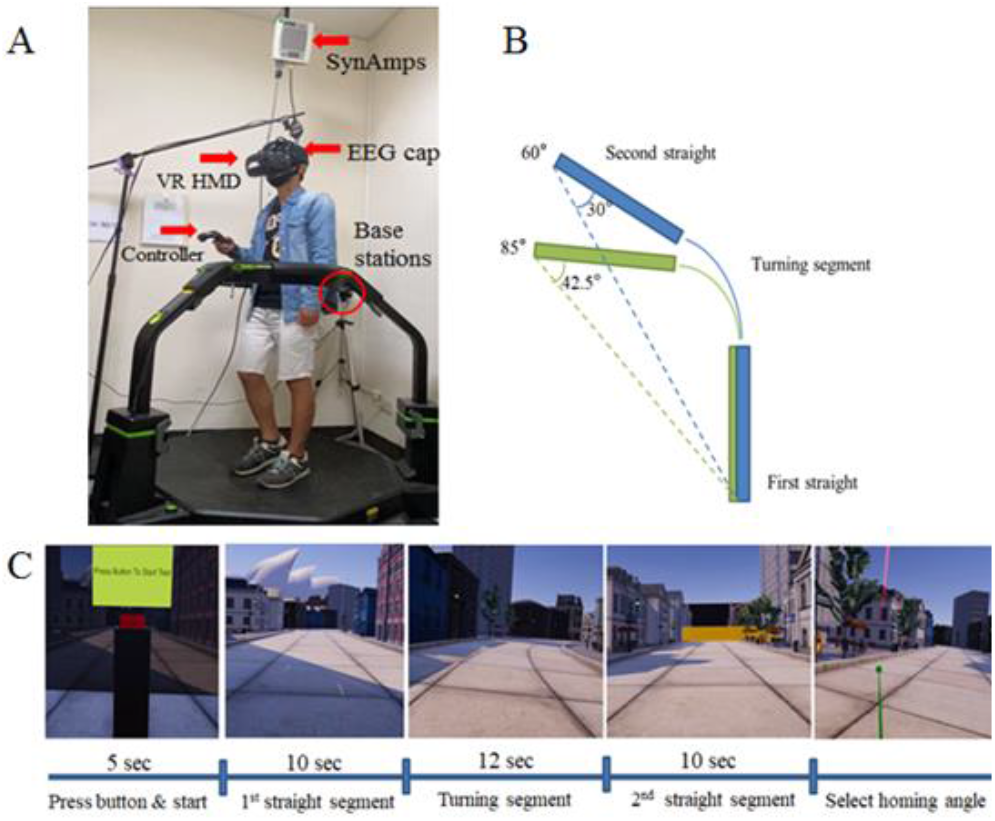
Experimental design of navigation trials. (A) A photo shows how a participant performed an experimental trial on the treadmill with a controller being held in his dominant hand. (B) A figure shows two paths with different turning angles. The 60° turn corresponds to a 30° homing angle, and the 85° turn corresponds to a 42.5° homing angle. (C) Screenshots of five segments in a trial. Participants pressed the red button, waited 5 seconds, and then started a trial. One turning path and two straight paths were presented before and after turning. Participants needed to keep turning along the path and keep themselves in the middle of the street. Upon arriving at the end point, participants were to turn back to select the homing angle with the controller, shown as the red arrow in the last screenshot.

Our VR walking scenario was made using Unity and based on snapshots of streets in Sydney and Opera House. In our active navigation tasks in the 3D scenario, there were five segments in a trial (Fig. 3C). Participants pressed the bottom area in front of the starting point, waited 5 seconds, and then started a trial. Participants were then presented with the first straight segment, then a turn, and then the second straight segment. After reaching the end point, they had to turn their body to select the homing angle with the HTC controller as the red arrow in the VR scenario. A trial was complete once participants selected the homing angles and then selected the ‘finish’ key; then, participants started the next trial. During the task, participants were asked to keep walking along the path in the middle of the street. The 1st and 2nd straight segments both took approximately 10 seconds to finish, and the turning part took approximately 12 seconds. In this task, we had two turning angles, 60° and 85°, which corresponded to 30° and 42.5° homing angles, respectively. (See Fig. 3B.) Two different turning-angle paths occurred randomly, and turning left or right was also random. Participants needed to finish 36 trials, including 18 trials for each homing angle with a 5- to 10-min break between half numbers of trials.

## III. data analysis

### A. Behavior Data Analysis

All homing data were categorized as nonturners with 42.5° and 30° homing-angle paths and turners with 42.5° and 30° homing-angle paths. There were a total of 1542 homing data points. After eliminating extreme values with deviations more significant than 25°, 48 data points were abandoned. Over 96.8% of the homing data were evaluated in statistical analysis. We calculated the mean value in each group and compiled the homing error chart with standard deviations using the Excel tool function, STDEV. The last step was to compute the significance between each pair of groups. Independent t-tests in Excel were used for the behavioral data (homing angle) analysis (comparison of accuracy). We compared the overall results of both strategy groups and different angle paths to examine whether the experimental procedure reached our expectations and resulted in a comparable accuracy level.

### B. Raw EEG Data Processing and Analysis

All EEG data were analyzed by the tools MATLAB and EEGLAB [23]. Thirty-two-channel EEG signals were first downsampled from 500 Hz to 250 Hz and then filtered to remove frequency components above 45 Hz and below 1 Hz to remove line noise (60 Hz and its harmonics) and DC drifting. Artifacts contaminated in the EEG signals were first identified by visual inspection using the EEGLAB visualization tool and eliminated to enhance the signal-to-noise ratio. Then, we used the EEGLAB plugin clean_rawdata(), a suite of EEG data preprocessing methods including artifact subspace reconstruction (ASR) [23, 24], which plays a central role in correcting continuous data [25]. After removing artifacts, adaptive mixture independent component analysis (AMICA), which was developed by Jason Palmer [26], was applied to the EEG data to extract independent components (ICs) from scalp electrode signals reflecting maximally statistically independent source time series. Then, event-related epochs were extracted from each trial and aligned for further time-frequency analysis. Each experimental trial consisted of one turning segment (12 seconds) and straight segments (10 seconds) before and after turning, and the first straight segment was used as the baseline for each segment.

All ICs with a residual variance of the equivalent dipole model of less than 15% were clustered based on measures including the time course of event-related potentials, mean IC log spectra, equivalent dipole locations, and event-related spectral perturbations (ERSPs) [15, 27]. First, we clustered components with topoplots by eye and then use the tool named ‘Talairach Client’[28, 29] to check the nearest gray matter of dipole location. In this way, we could double check whether the component we selected suited the component cluster. After clustering the ICs near the cortices, we wanted to see the results for brain regions, including the parietal, occipital, frontal, and central regions, and then we plotted the ERSPs for nonturners and turners with each angle path and the difference in ERSPs between the two strategies [4–6].

## IV. results

### A. Behavior Results

The mean homing responses of all active and passive navigation participants are displayed in Fig. 5 for both turners and nonturners indicated as different color lines. The 60° and 85° turning-angle paths corresponded to 30° and 42.5° homing angles, respectively. In active navigation tasks (Fig. 5A and 5B), turners revealed higher accuracy than nonturners for low eccentricity (30°) with significant difference *(p* < .001); in contrast, nonturners were more accurate at higher eccentricity (42.5°) without significant difference *(p* = .126). Turners overestimated the homing angle in small turning-angle paths, whereas they underestimated large turning-angle paths, and nonturners underestimated homing angles in both angle paths. In passive navigation tasks (Fig. 5C and 5D), turners revealed higher accuracy than nonturners at both low (30°) eccentricity with significant difference (*p* < .05) and high (42.5°) eccentricity without significant difference *(p* = .110). All mean angles that nonturners and turners selected were underestimated in both 42.5° and 30° homing-angle paths.

**Fig. 4.**
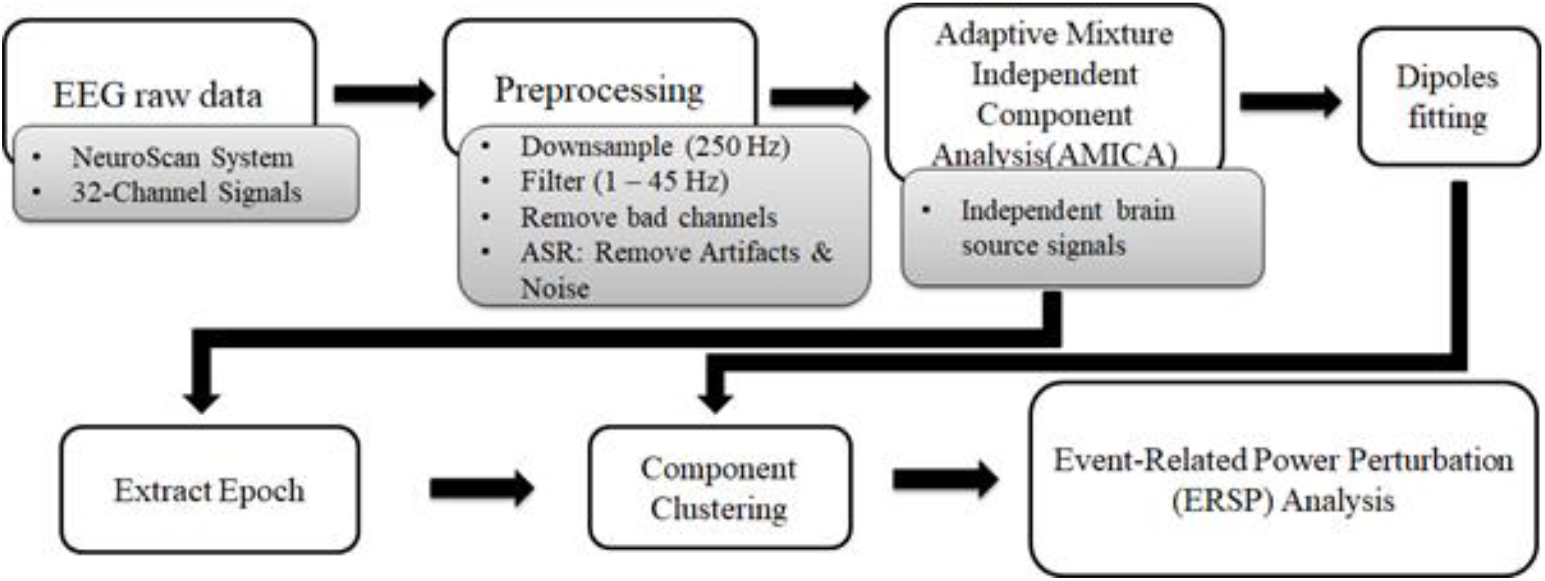
Flow chart of EEG data processing. Raw EEG data were first preprocessed, including downsampling, filtering, and artifact removal; then, we applied AMICA to independent brain sources. In the next step, we ran dipole fitting to obtain the 3D positions of the brain sources. Epochs were extracted for three segments: the 1st straight, turning, and 2nd straight segments. After checking the topoplot and dipole position, we clustered all the components located in or near the cortex we wanted and then plotted the ERSPs for each brain cortex.

**Fig. 5.**
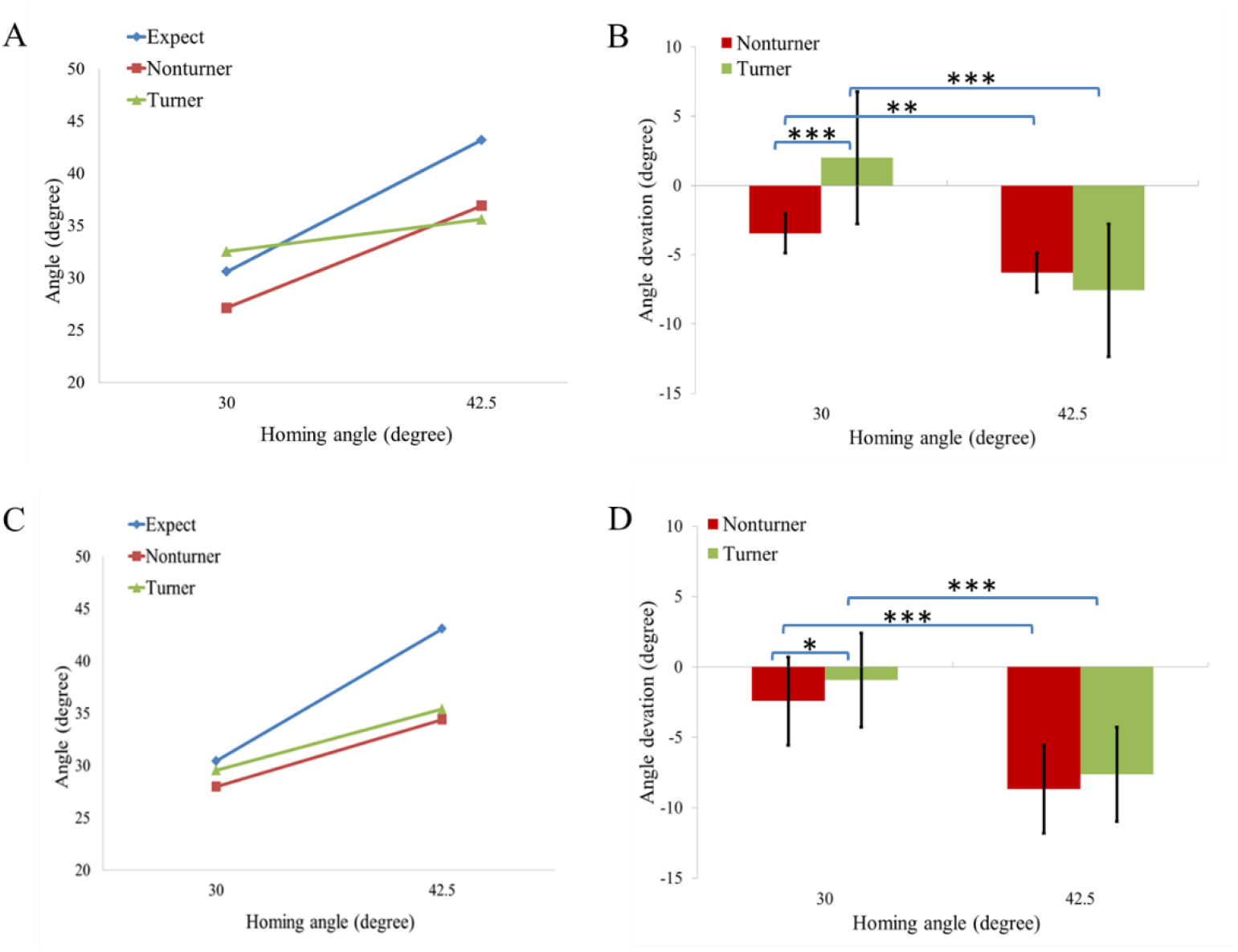
Homing performance of turners and nonturners. (A) Homing angle reported by participants (Y-axis) as a function of homing angle (X-axis) in the active tasks. Blue line represents the expected homing angle. Red and green lines represent the homing angle chosen by nonturners and turners, respectively. (B) Signed homing error as a function of homing angle in active tasks. Error bars indicate standard error. *: *p* < .05, **: *p* < .01, ***: *p* < .001. (C) Homing angle reported by participants in passive tasks. (D) Homing error of participants in passive tasks.

A 2 x 2 x 2 factorial mixed-measure ANOVA was conducted to examine the effects of spatial reference frames (egocentric and allocentric) and experimental environments (active and passive walking) on expected homing angles, homing responses, and deviations.

Due to the slight differences in location when participants conducted the pointing task, the expected homing angles had nuances compared to designed homing angles (30° and 42.5°) among different participants. The ANOVA results revealed a statistically significant difference between the two angle types, F(1, 736) = 321206.465, *p* < .000, partial n^2^ = .997. Besides, there were no statistically significant interactions between either spatial reference frames or experimental environments on homing angles. For spatial reference frames on homing angles, the interaction result was F(1, 736) = .075, *p* = .785, partial n^2^ = .0001; for experimental environments on homing angles, the interaction result was F(1, 736) = .179, *p* = .673, partial n^2^ = .0002; for spatial reference frames and experimental environments on homing angles, the interaction result was F(1, 736) = 3.748, *p* = .053, partial n^2^ = .005.

Concerning homing responses, the main effect of angle types (30° and 42.5°) was significant, F(1, 736) = 410.343, *p* < .000, partial n^2^ = .358. There was a statistically significant interaction between spatial reference frames and experimental environments on responses, F(1, 736) = 23.249, *p* < .000, partial n^2^ = .031. Therefore, an analysis of simple main effects for experimental environments was performed with statistical significance with a Bonferroni adjustment and was considered significant at the *p* < .025 level.

The mean homing response to 30° for egocentric walking in an active environment (M = 32.243, SE = .587) was significantly higher than that for walking in a passive environment (M = 29.556, SE = .590), at 2.687 (95% CI, 1.052 to 4.322), *p* = .001. In addition, when walking in an active environment, the mean homing response to 30° for the allocentric participants (M = 26.861, SE = .569) was significantly lower than that for the egocentric participants (M = 32.243, SE = .587) by 5.383 (95% CI, 3.777 to 6.989), *p* < .000.

For mean homing responses to 42.5°, there was a statistically significant difference between active and passive environments for allocentric participants. The mean homing response to 42.5° with active walking (M = 36.900, SE = .687) was significantly higher than that with passive walking (M = 34.406, SE = .736), 2.495 (95% CI, .518 to 4.471), *p* = .013.

Additionally, there was also a statistically significant interaction between spatial reference frames and homing angle for responses (30° and 42.5°), F(1, 736) = 36.088, *p* < .000, partial n^2^ = .047. Simple main effects analysis revealed a statistically significant difference in homing responses to 30° between the egocentric and allocentric participants, F(1, 747) = 34.522, *p* < .000.

Concerning homing deviations, the main effect of angle types (30° and 42.5°) was significant, F(1, 736) = 200.147, *p* < .000, partial n^2^ = .214. There was also a statistically significant interaction between experimental environments (active and passive walking) and homing angle for deviations (30° and 42.5°), F(1, 736) = 10.442, *p* = .001, partial n^2^ = .013989. Simple main effects analysis revealed that there were statistically significant differences in homing deviations to both 30° and 42.5° between the egocentric and allocentric participants, F(1, 747) = 80.490, *p* < .000 at 30° and F(1, 737) = 9.034, *p* = .003 at 42.5°.

### B. EEG Results

Mean ERSP images during navigation in parietal, occipital, frontal, and central clusters for turners and nonturners and their differences are shown in Fig. 6(A)-(D).

**Fig. 6.**
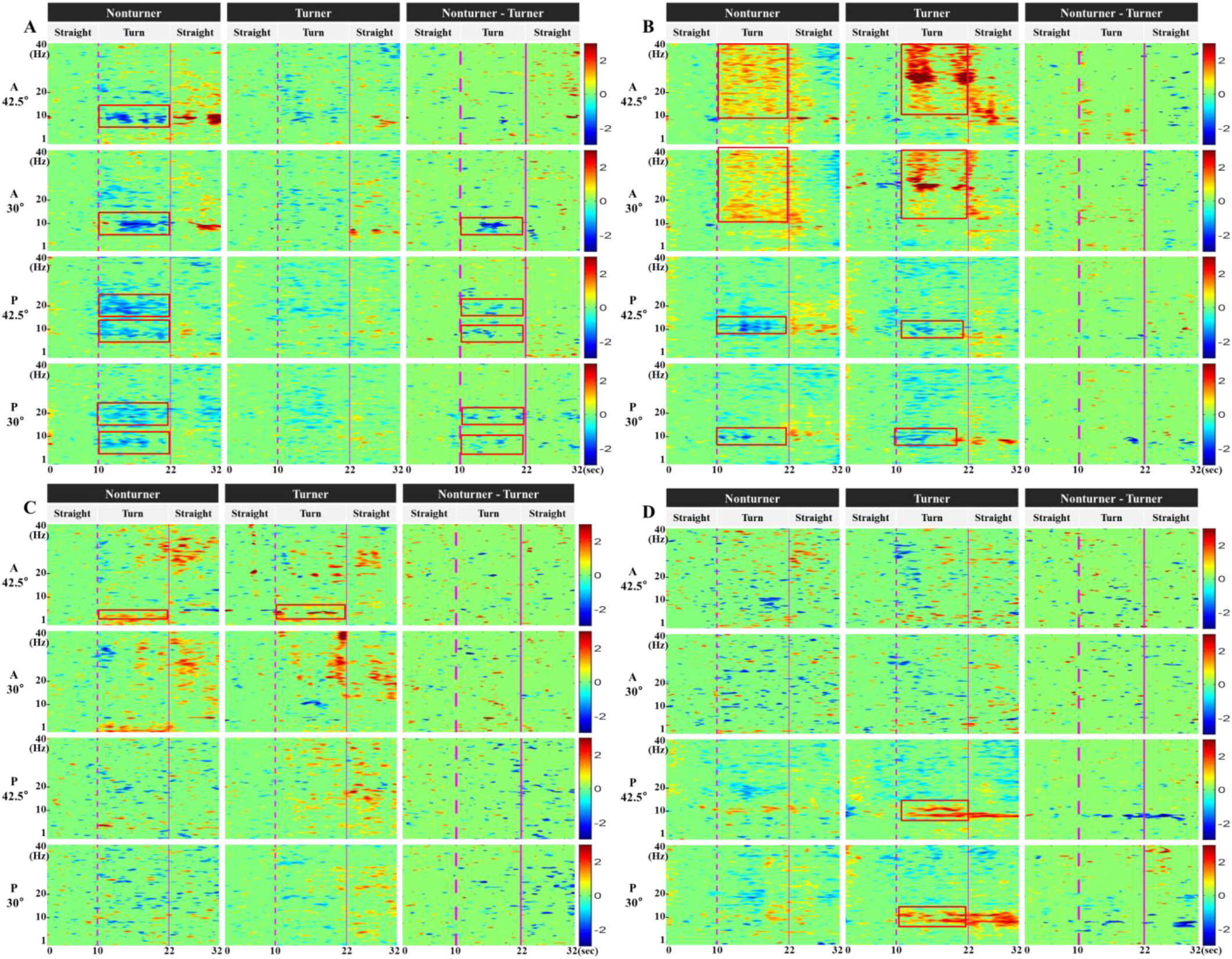
Event-related spectral perturbation (ERSP) for selected clusters. Leftmost numbers and characters show the homing angle and the active (A) or passive (P) task for the ERSP figures with frequencies from 1 to 40 Hz. The first column displays nonturner ERSPs during navigation paths. The middle column displays turner ERSPs during navigation paths. The right column displays the difference between nonturners and turners. (A)-(D) Mean ERSP images during navigation in the parietal (2A), occipital (2B), frontal (2C) and central (2D) clusters for turners and nonturners and their differences.

In the parietal cortex (Fig. 6A), we observed that nonturners demonstrated alpha desynchronization, accompanied by a decrease in alpha-band activity during large and small-turning tasks. In addition, turners had no obvious power change feature in ERSPs for either navigation task. However, when we compared two strategies by subtracting nonturner ERSPs from turner ERSPs, the alpha difference still remained. This result demonstrated that these two strategies of participants in large angle tasks had more similar power-changing features in the parietal alpha band. In the occipital cortex (Fig. 6B), both turners and nonturners showed power increases in beta and low gamma bands while turning. Turners had even, more substantial power enhancement in high beta and low gamma frequency than nonturners.

Moreover, large-angle paths caused more obvious power increases in high-frequency bands in turners. Regarding the frontal ERSP results (Fig. 6C), both nonturners and turners showed dominant power increases during large turning tasks. Small turning task ERSPs did not reveal significant theta power changes for either nonturners or turners. Central cluster ERSPs (Fig. 6D) revealed only fragmentary and small patches of power decreases during both large and small turning tasks for nonturners and turners. We could barely see any significant features in the central cluster ERSPs.

### C. Correlation with the Parietal Cortex

As we know from previous studies that the main brain dynamic features of people performing navigating occur in the parietal cortex in alpha and beta bands [4–6], we computed the correlation R of brain power between the parietal cortex and the occipital cortex, between the parietal cortex and the frontal cortex and between the parietal cortex and the central cortex to reveal the activity of these cortices during navigation tasks. Table 1 demonstrates the correlation results in both active and passive navigation tasks. We used the mean power of the occipital, frontal, and central mean ERSPs at every single time point during turns in each frequency band to determine the correlation with the parietal cortex.

**TABLE I.** Correlation Results

| Cortex \ Frequency band | Nonturner 42.5 |  |  | Nonturner 30 |  |  | Turner 42.5 |  |  | Turner 30 |  |  |
| --- | --- | --- | --- | --- | --- | --- | --- | --- | --- | --- | --- | --- |
|  | Theta | Alpha | Beta | Theta | Alpha | Beta | Theta | Alpha | Beta | Theta | Alpha | Beta |
| Occipital | -0.0462 | 0.061 | 0.1765 | -0.2084 | -0.1121 | -0.1164 | 0.345 | -0.3439 | -0.1469 | -0.125 | -0.2878 | 0.2033 |
|  |  |  |  |  |  |  | * | * |  |  | * |  |
| Frontal | 0.57 | 0.4228 | 0.5357 | 0.2264 | -0.1942 | 0.7282 | -0.2539 | 0.7844 | 0.3097 | 0.4396 | -0.0404 | 0.2342 |
|  | *** | *** | *** | * |  | *** | * | *** | ** | *** |  | * |
| Central | 0.5884 | 0.3243 | 0.3047 | -0.2807 | 0.05 | 0.5889 | 0.0664 | 0.699 | 0.8967 | 0.2918 | 0.0293 | 0.5286 |
|  | *** | ** | ** | ** |  | *** |  | *** | *** | ** |  | *** |

| Cortex \ Frequency band | Nonturner 42.5 |  |  | Nonturner 30 |  |  | Turner 42.5 |  |  | Turner 30 |  |  |
| --- | --- | --- | --- | --- | --- | --- | --- | --- | --- | --- | --- | --- |
|  | Theta | Alpha | Beta | Theta | Alpha | Beta | Theta | Alpha | Beta | Theta | Alpha | Beta |
| Occipital | 0.3175 | 0.0445 | 0.5766 | 0.5717 | 0.4829 | 0.1706 | 0.076 | -0.1036 | 0.4611 | 0.0237 | 0.8336 | 0.6599 |
|  | ** |  | *** | *** | *** |  |  |  | *** |  | *** | *** |
| Frontal | -0.0425 | -0.0014 | 0.207 | -0.0197 | 0.8211 | 0.3296 | -0.3139 | 0.4734 | -0.1238 | 0.1083 | 0.4528 | 0.1956 |
|  |  |  |  |  | *** | ** | ** | *** |  |  | *** |  |
| Central | 0.2565 | 0.5789 | 0.6955 | 0.3472 | 0.7607 | 0.9215 | -0.3569 | -0.0538 | -0.2162 | 0.3717 | 0.4903 | 0.7943 |
|  | ** | *** | *** | ** | *** | *** | ** |  |  | *** | *** | *** |

Columns are names of the cortices, rows are nonturners and turners in 42.5° and 30° homing-angle paths in different frequency bands, including theta (4 – 7 Hz), alpha (8 – 14 Hz) and beta (15 – 30 Hz) bands. \*: $p < .05$ , \*\*: $p < .01$ , \*\*\*: $p < .001$ .

## V. Discussion

In this study, we investigated the differences in brain activation during active and passive navigation tasks and the homing angle data after navigation between participants using allocentric and egocentric spatial reference frames. As per pret-test, we already knew which spatial strategy the participants preferred to use. However, the RFP was not permanent. Participants may have preferred a different reference frame in a different environment. For example, participants in the passive environment (pretest) in this study, although they could perceive structured allocentric information, may have changed their RFP in active navigation tasks (a scenario in VR).

Passive navigation tasks were successfully observed in navigation-related neural mechanisms [4, 8]. However, several components strongly contribute to active navigation, including proprioception, vestibular input, motor activation, and attention and planning allocation [13, 30–32]. Thus, we have investigated if more brain areas and interactions between brain areas may be involved in active navigation tasks than passive navigation tasks.

## A. Homing Angle with Different Turning Angles and Reference Frames

In the mean homing angle result shown in Fig. 3, we can see that only turners in small turning trials tend to overestimate. Turners in large turning trials and nonturners in different turning trials underestimated the homing angle related to the start point. This homing angle result is quite similar to previous passive navigation results [6]. Second, after we checked each strategy’s standard deviation in the homing error results (Fig. 4), turners revealed a higher deviation than nonturners. This finding could indicate that nonturners have more general spatial navigation abilities and are better navigators than turners [33]. Homing errors from both turners and nonturners in different turning-angle trials had significant differences, as did two different reference frames in small turning trials. Only two different types of participants had no significant difference in large turning trials. It seems that while participants applied specific spatial RFPs in the large turning-angle trials, the performance of allocentric and egocentric reference frames tends to be similar. In the next part, we investigated this outcome from the perspective of EEG dynamics in widespread brain regions.

## B. EEG Dynamics with Distinct Reference Frames in Spatial Navigation

Previous studies have suggested that the use of egocentric and allocentric reference frames activates widespread overlapping cortical networks. In this study, common brain regions, including the frontal, central, parietal, and occipital cortex, were explored during active navigation for both turners and nonturners using an egocentric or an allocentric reference frame, respectively. Our successful navigation involved significant EEG modulation power changes that were dominant in the theta (4-7 Hz), alpha (8-14 Hz), and beta (15-30 Hz) frequency bands between widespread brain regions.

As a multisensory area, the parietal cortex is known to support the integration of information from different sensory modalities. It is based on distinct spatial coordinate systems. Parietal alpha power modulation was previously observed during spatial exploration and orientation maintenance [4, 16, 17]. Alpha desynchronization in or near the parietal cortex was significantly stronger when approaching and during the turn in nonturners than in turners in passive navigation [4]. In passive navigation, as a control group, nonturners also revealed power suppression in alpha and beta frequency bands (Fig. 5A). However, in active navigation, allocentric participants may have changed their previously determined preference, and participants responded in an egocentric reference frame while selecting homing angles [13]. The parietal component cluster exhibited significantly stronger alpha-blocking in both large and small turning-angle tasks in our navigation task. Parietal ERSPs did not reveal significant differences, whereas large turns did reveal significant differences between nonturners and turners. Then, we compared previous study ERSP results in the parietal cortex [4]. The results showed even more apparent and stronger alpha-blocking for nonturners. Therefore, we considered that allocentric participants might change their spatial RFP and become egocentric navigators while they are in an active environment, and a large-angle path presented to allocentric participant which may cause them to have a tendency to switch into an egocentric reference frame.

The occipital cortex also plays a vital role in navigation tasks. Most previous studies showed that participants demonstrated power modulation in the alpha and beta bands [4, 6, 13], and our passive results also demonstrated the same features (Fig. 5B). By contrast, our occipital ERSP results of active navigation showed no features in the alpha and low beta bands. Instead, power increased in high beta and gamma frequency bands while turning for both nonturners and turners. Turners’ high-frequency power increase seemed to be stronger than that of nonturners in the respective occipital ERSPs. According to Catherine [34], “In humans, scalp EEG recordings consistently reveal the existence of synchronized oscillatory activity in the gamma range when participants experience a coherent visual percept”. Studies of working memory load effects on human EEG power have indicated that gamma power typically increases with load [35–37]. Moreover, the results from M. Gola et al. [38] showed that beta activity in occipital regions is perfectly correlated with an attentive visual performance. These studies seem to be the basis for our occipital cluster ERSPs. In active navigation, participants were to turn to follow the course of the path. The heading was consistent with the orientation, leading to coherent visual perception and resulting in beta and low gamma power synchronization. Turners needed to update their position and orientation in their minds, though nonturners only updated position information. This might be why turners revealed more substantial power increases than nonturners in the high-frequency bands. In the same way, turners also demonstrated stronger high-frequency activity in larger-angle tasks than in small-angle tasks. However, we could not rule out the possible impact of artifacts from muscle movement. Because participants should turn their body and neck while turning, muscle movement may have an impact on EEG channels near the occipital cortex, including O1, O2, and Oz, and lead to a power increase in the high-frequency band. We ran the correlation of mean power between occipital and parietal regions in each frequency band to verify this hypothesis as we know that the parietal cortex has a primary role in navigation, regardless of previous studies or our own results. As a result, all four groups of homing angle data in the beta frequency band did not reveal significant correlations (*p* > .05). This could be evidence that the power increase in the occipital cortex was not caused by navigation.

Previous studies revealed that the frontal-parietal network is for resolving allocentric and egocentric reference frames [39, 40]. The ACC is assumed to underlie spatial learning, visual attention, orientation, working load memory, and path planning [41, 42]. In our active navigation task, a cluster in or near the frontal cortex demonstrated significant theta power synchronization in large-angle turning, likely reflecting spatial working memory demands for successful path integration (Fig. 5C). Accurate homing responses in this task required participants to update the starting location relative to their end position. Substantial theta power increases during turning reflect increasing task requirements related to upcoming heading changes. This finding replicates previous statements of increased theta power in the frontal region while performing more demanding spatial navigation tasks [4, 43, 44].

In the large-angle trials, turners that used egocentric references demonstrated more pronounced theta activity than nonturners that used an allocentric reference frame. This is in light of the higher amount of information that has to be updated within an egocentric reference frame, such as orientation and position changes with each rotation and transformation. In contrast, an allocentric reference frame only requires updating of position rather than the orientation of the navigator [1]. In conclusion, egocentric spatial updating requires a greater working memory load than the use of an allocentric reference frame for spatial updating [42, 45]. However, both turners and nonturners had no theta power increase in the frontal cluster in the small turning trials. In our small turning-angle scenario, participants could see the end point soon after the turning started, which meant that participants did not need to keep updating their position and orientation because they already knew the end point was there. Less spatial learning would result in less power increase in the theta frequency band in the frontal cluster. In the ERSP results of passive navigation, we did not observe obvious power disturbances. The probable reason might be that participants would drift off or be distracted easily while watching the screen in front of the table rather than wearing the VR head-mounted display in active navigation.

Central cluster ERSPs of active navigation revealed only fragmentary and small patches of power changes (Fig. 5D) during both large and small turning tasks for nonturners and turners. However, they could not be regarded as significant features while navigating. Previous passive navigation results revealed alpha modulation in motor and central regions and assumed alpha suppression to reflect motor cortex activity during imagined movement [6]. Motor imagery can secure brain oscillations in Rolandic mu rhythm and central beta rhythm, originating in the sensorimotor cortex [46]. We also found that nonturners in passive navigation have the same feature in central cluster ERSPs. In our active navigation task, participants were actually moving their bodies during trials. Therefore, conceivably, it is possible that the central cluster ERSPs did not show significant features within the two strategies used by the participants. In passive ERSP results in the central cortex, the increase in turners’ activity near 10 Hz during turns was not reported clearly, but it might relate to motor imagery or saccadic and optokinetic eye movements during visual flow stimulation [4, 47–49].

## VI. Future works and limitations

As we found the phenomenon that people will switch spatial strategies when in a different environment in our active experiment, we believe the experimental procedure proposed in this paper could also be used to investigate more important and further questions like which factors cause people to have a tendency to change spatial reference frames. With direct control of the switching strategy, we can guide people to use an appropriate and efficient strategy for a navigation task and train them to be more sensitive to their environment. Furthermore, we can expect that the sense of direction with the use of brain dynamics can be applied to spatial learning, real-world driving, detour management, and even rescue training. In future, following could be reduce as well for better results:

1. Since our experiment included active navigation, more noise and artifacts were caused by body movements in EEG signals than in passive experiments. We could not ensure the elimination of all noise and artifacts. Therefore, a better EEG recording sensor or artifact eliminator to preclude all noise and muscle movement impact would be another essential aspect to consider.
2. During the experiment, a wireless EEG recording system was better for an active experiment. Furthermore, we manually adjusted the belt of the HMD to avoid contact with the sensors on the EEG cap. This might not have been possible if caps with higher sensor densities were used. We believe the integration of the EEG cap with the HMD is natural, and we expect to see commercial products from companies such as MindMaze to be available on the market soon.
3. Our current setup used the Scan 4.5 system, and the recorded EEGs were analyzed offline. This device is only suitable for an initial investigation in a lab environment due to its long setup time. We believe it should be possible to reproduce the results using off-the-shelf, portable EEG devices and to process the data in real-time.

## VII. Conclusion

In this study, we investigated the general spatial navigation abilities and brain dynamics between the preference of egocentric and allocentric frames of reference in an active virtual navigation task. In this VR navigation environment, we indeed observed some differences in RFP. Behavior results revealed that nonturners were better than turners in homing navigation performance. We also observed strategy-dependent brain dynamics for spatial navigation and the evidence that strategy will change when people are in different situations. We observed the power changes and causal flow patterns in ERSPs in parietal occipital and frontal cortices with different RFPs and different angle paths. Our behavior results are consistent with parietal ERSP results and strongly support the assumption that people will switch their spatial reference frame in different environments, rather than the assumption that only one spatial reference frame is used to solve a task. However, since a good navigator should be able to identify the appropriate strategy for a navigation task, the factor and the direct control of the switching strategy are very important for us to improve the human capability of navigation.

## Footnotes

1 https://www.virtuix.com

